# Spatiotemporally distinctive astrocytic and neuronal responses to repetitive intracortical microstimulation

**DOI:** 10.64898/2026.01.02.697363

**Authors:** David Grundfest, Kayeon Kim, Alexandra Katherine Isis Yonza, Jeremi Tadeusz Podsiadlo, Lechan Tao, Xiao Zhang, Krzysztof Kucharz, Yan Zhang, Anpan Han, Barbara Lind, Changsi Cai

## Abstract

Astrocytes are increasingly recognized as active modulators of neuronal synaptic transmission. Intracortical microstimulation (ICMS) is widely used to manipulate neuronal activity, yet the accompanying astrocytic responses remain poorly characterized. Using dual-color *in vivo* two-photon calcium imaging to simultaneously monitor neurons and astrocytes, we show that ICMS elicits astrocytic activation with spatiotemporal features that diverge from those of neurons. Astrocytes were recruited at stimulation intensities as low as 10μA, thresholds sufficient to activate neurons, indicating that astrocytes robustly sense electrical perturbation. Unlike neurons, however, astrocytic responses were spatially heterogeneous and temporally variable across trials. At higher stimulation intensities (>=50μA), astrocytic responsiveness, i.e., response peak amplitude, and number of responsive trials, progressively attenuated across repeated trials, in contrast to the stable and consistent neuronal responses. Although neuronally driven, astrocytes exhibited a distinct response profile under the same stimulation parameter, revealing a unique component of electrically evoked cortical activity that underscores the importance of incorporating glial physiology into future neuroprosthetic strategies.

## Introduction

Neurons and astrocytes are integral, interacting components of cortical circuits [1,2]. While neurons encode and transmit information [3–5], it is increasingly appreciated that astrocytes play an active role in shaping neuronal signaling. Beyond their classical functions in ionic and metabolic homeostasis [6–8], astrocytes participate in neurotransmission by releasing gliotransmitters, and modulate synaptic efficacy [9]. Astrocytic Ca^2+^ elevation occurs not only in somata but also within highly localized microdomains in fine processes, which are spatially restricted and temporally diverse [10,11]. These microdomains respond to local synaptic glutamate release, neuromodulators, and local circuit activity [12], placing astrocytes in a position to modulate synaptic transmission and plasticity while coordinating local circuit states [13,14]. *In vivo* sensory stimulation studies further show that astrocytic microdomains are activated on timescales comparable to neuronal responses, revealing substantial functional heterogeneity and regional specificity [15,16]. Such spatially compartmentalized Ca^2+^ signals highlight astrocytes as distinct functional units within cortical circuits [17], capable of tuning neuronal activity and contributing to information processing in a manner that extends beyond passive support.

Intracortical microstimulation (ICMS) is widely used to probe and modulate cortical function in both basic research and neuroprosthetic applications. Its clinical relevance has expanded rapidly with the use of ICMS in humans for brain-machine interface systems [18–22]. However, despite this growing adoption, most ICMS studies have focused on neuronal responses, leaving astrocytic contributions in this context far less understood. Previous studies have provided multidimensional characterization of how ICMS engages cortical neural circuits. These works show that the spatial extent of neuronal recruitment increases in a non-radial pattern with stimulation intensity, with activation probabilities determined largely by axonal geometry, proximity to the electrode, and the distributed patterns of axonal activation [23–25]. Temporal response patterns vary systematically with pulse width, frequency, and train duration, demonstrating distinct temporal firing regimes and frequency-dependent adaptation [26,27]. Cell-type-specific analyses further show that excitatory and inhibitory neurons differ in their activation thresholds and response persistence [28,29]. Although electrical stimulation paradigms have been used to evoke astrocytic activity [30–34] and to study neurovascular coupling (NVC) [35,36], astrocytic responses to ICMS and their interaction with neuronal activity remain poorly understood. Previous studies have reported limited astrocytic Ca^2+^ signals in response to ICMS [23], noting in particular that astrocytes are difficult to recruit at lower stimulation intensities *in vivo* using high frequency, and prolonged stimulation durations [27].

Importantly, these earlier calcium-imaging studies relied on sulforhodamine 101 (SR101) labeling and Oregon Green BAPTA (OGB) indicators, method that provided bulk astrocytic signals [37] but lacked the resolution needed to resolve fine response dynamics, often limiting analyses to somatic activity. This method likely overlooked microdomain events, now recognized as a key locus of astrocytic signaling and cortical computation with greater complexity [32,38], and fundamental to neuron-astrocyte communication [39–41]. In contrast, in the current study, we use a GFAP promoter to drive expression of a membrane-anchored variant of the Ca^2+^ indicator GCaMP6f to monitor astrocytic Ca^2+^ transients. This approach provides high-fidelity detection of astrocytic activity across both somatic and microdomain compartments [42–44]. This methodological improvement allowed us to directly compare astrocytic and neuronal calcium dynamics under ICMS with single-cell resolution, raising the possibility that astrocytic microdomains may be important sites of ICMS engagement.

A central gap is the lack of systematic, cellular-resolution characterizations of astrocyte versus neuron spatiotemporal responses to ICMS across defined current intensities and repeated trials. It is unclear (1) whether astrocytes can be reliably recruited at lower currents commonly used in neurophysiology and neuroprosthetics, (2) how astrocytic response amplitude compare with co-localized neuronal dynamics and (3) how spatial factors-such as soma or region of interest (ROI) -to-electrode distance- and temporal dynamics differentiate neuronal and astrocytic recruitment. Clarifying these issues is essential for interpreting ICMS-evoked signals, including hemodynamic readouts, and for designing stimulation strategies that either leverage or evade astrocyte-mediated effects.

To address these questions, we combined dual-color two-photon calcium imaging with co-labeled astrocytic and neuronal indicators in mouse visual cortex, allowing direct comparison of astrocytic and neuronal responses under identical ICMS parameters. Astrocytic ROIs were defined using a state-of-the art grid-based analysis approach [45] that captured both somata and fine-process signals, enabling detection of microdomain-level activity without separating compartments. Through detailed resolution of astrocytic dynamics in an unbiased way, this approach establishes a systematic characterization of their spatiotemporal responses together with neuronal activity.

## Materials and Methods

### Animals

Experiments were performed on C57BL6 mice (N=5). All animal procedures complied with Directive 2010/63/EU of the European Parliament and the Council on the protection of animals used for scientific purposes. Protocols were reviewed and approved by the Danish National Committee on Health Research Ethics, in accordance with the European Council’s Convention for the Protection of Vertebrate Animals used in experimental and other scientific studies.

### Experimental procedures

#### Viral vector injection

Two weeks prior to the recording sessions, animals received intracortical injections of a viral mixture (9:1 ratio) targeting two sites within the visual cortex (+1mm AP, + 2mm ML relative to lambda) with two depths (2×300nl). The mixture consisted of AAV5-gfaABC1D-Ick-GCaMP6f, which labels astrocytic calcium dynamics, and rAAV-CaMKIIa-jRGECO1a-WPRE-hGH polyA (titer: 2.48E+12vg/m), which selectively expresses a red fluorescent calcium indicator in excitatory neurons (Fig. 1A, B).

**Figure 1.**
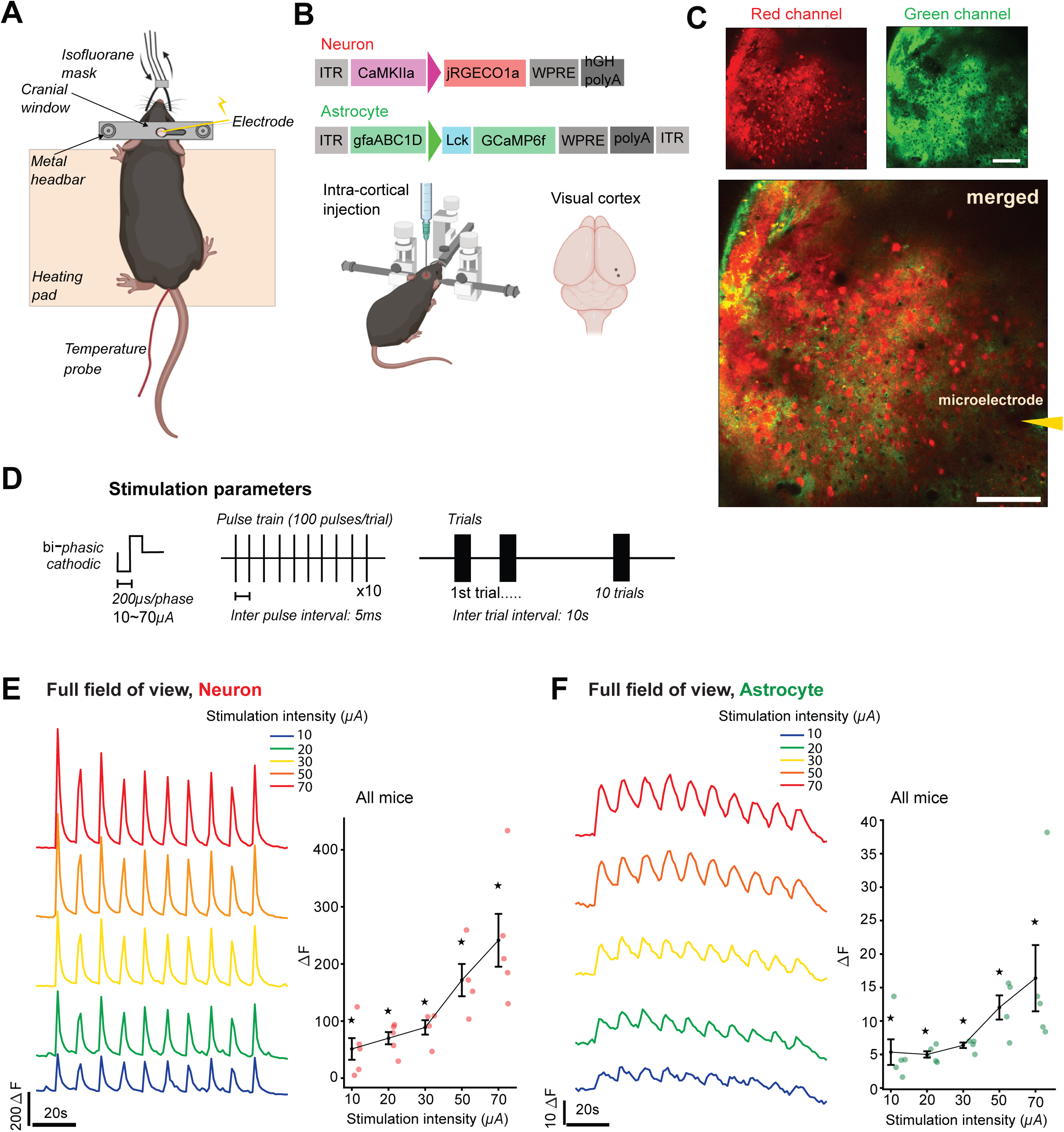
Experimental overview and robust recruitment of neurons and astrocytes by low-intensity ICMS. **A.** Schematic of the experimental setup: a microelectrode was inserted into the visual cortex of a mouse. **B**. Diagram of the viral vectors. A mixture of viral constructs was injected intracortically at two sites within the visual cortex (see *Methods* for details). **C**. Representative two-photon images showing a co-labeled neuron (red, upper left), astrocyte (green, upper right) and the merged image (bottom). Scale bar, 100µm. **D**. Stimulation waveform and parameters (see *Methods* for details). **E**. Example neuronal response amplitudes across stimulation intensities during ten repetitive stimulations from a single mouse. Left: averaged traces across the whole field of view. Right: group-averaged response amplitudes plotted as a function of stimulation intensity. Each dot represents one animal; error bars indicate mean ±SEM. \**p*<0.05, one-sample *t*-test against zero. **F**. Same as in (E) but showing astrocytic response amplitudes.

#### Stimulators and stimulation protocol

We employed ISO-flex (A.M.P.I) connected to platinum-iridium micro-electrode (8-11kΩ, tip diameter, 2-3µm; PI2PT30.01. A3; Microprobes for Life Sciences). A return electrode was placed subcutaneously under the mouse’s neck. The stimulating electrode was mounted in a micromanipulator arm (Fig. 1A) at an insertion angle of 15-20° for two-photon imaging and positioned 200µm below the cortical surface to set the stimulation depth. Electrical stimulation consisted of cathodic-leading biphasic pulses (200 µs per phase, without interleaving between pulses; Fig. 1D), with stimulation duration of 500ms. Stimulation was delivered at a fixed frequency of 200 Hz across all sessions and animals. Each experiment included 5-10 stimulation trials, with a 10-second inter-trial interval [46].

#### Animal surgery and preparation

The surgical procedure was carried out as previously described [46]. For the acute experiment, anesthesia was induced with isoflurane of 4%, and then maintained throughout the experiment at a lower concentration ranging from 0.9 to 1.5%. After checking animal reflexes, local anesthesia was first administered with lidocaine (10mg/kg) prior to making the surgical incision. After performing craniotomy, the dura mater was carefully removed to expose the cortical surface. Following this, the electrode probe was inserted into the cortex, and agarose gel was applied to both stabilize the cortical surface and secure the probe in place. To maintain physiological stability during the procedure throughout the experiment, the animal’s body temperature was carefully regulated at 37°C using a heating pad. Additionally, for pharmacological manipulation, Tetrodotoxin (TTX) was applied topically at a concentration of 20μM (Tocris Bioscience) to block neuronal activity.

### Two-photon imaging and data analysis

We performed imaging using a two-photon microscope (TPM) (FluoView FVMPE-RS, Olympus) equipped with a femtosecond laser (Mai-Tai DeepSee) and 25 × 1.05 NA water-immersion objective, with GaAsP detectors for signal detection. An excitation wavelength of 1000nm was used to detect both green and red fluorescence signals. Image acquisition rates ranged from 0.9 to 2.43Hz, adjusted according to the selected field of view and pixel resolution, which ranged from 0.994 to 1.989 µm/pixel.

#### Grid-based signal analysis

For each recording, a square grid was generated [45], with each square measuring ranging between 15 and 16 μm. Each square is considered a region of interest (ROI). A temporal signal coming from one ROI was computed as an average over all pixels within that square. From such a signal we calculated z-score using the average and standard deviation of that signal before onset of the first trial. Z-score signal was then low-pass filtered using moving average with window size 7. For filtered signals we calculated a baseline using asymmetric least squares with lambda = 30, *p* = 0.001 and 10 iterations. This baseline was subtracted from the signal to level the peaks so they could be thresholded. A ROI was classified as responsive to a stimulation trial if the signal of that ROI exceeded the threshold at value V and stayed above the threshold for a duration of D seconds within the specified time window. V was set to 2 for both neurons and astrocytes, D was set to 2 for both neurons and astrocytes. The time window for neurons was set to be 5 seconds immediately after stimulation onset for neurons and 5 seconds with 3 seconds delay for astrocytes. A ROI was considered responsive if it responded to at least one stimulation trial.

#### Analysis of response distribution shape

To quantify how stimulation intensity influences the shape of the distribution of neuronal and astrocytic responses, we computed a weighted average across the histogram of the number of responsive trials excluding bins with zero responses. Responsive number of trials from 1 to 10 were averaged, using the number of regions of interest (ROIs) in each bin as weights, thereby deriving the relative ratio among the number of responsive trials.

#### Segmentation of neuronal somata

Neuronal somata were segmented from an image calculated by averaging across time in the red neuronal channel. We then enhanced contrast by contrast stretching followed by median blur with kernel size 5. Blurred image was then thresholded by *adaptiveThreshold* function from opencv python library using Gaussian adaptive thresholding method with blocksize 15 and shift constant C = −15. To remove as many false positives as possible, the binary image was processed using a morphological opening with a cross kernel of size 3. Remaining positives were segmented to instance using watershed segmentation with a minimal distance 5. If a bounding box of an instance had an area bigger than 200 pixels, it was also considered false positive. All remaining segmentations were considered to be neuronal somata.

#### Calculation of neuronal soma occupancy in a grid

Neuronal somata were segmented and projected onto the astrocytic channel. For each ROI, soma occupancy was calculated as the percentage of area covered by somata. ROIs were classified as containing somata (=neuronal soma within the ROI) if occupancy exceeded 5% (Fig. 4F).

### Quantification and statistical testing

All statistical testing was performed using python library SciPy (http://scipy.org/ciring-scipy/). We used the Benjamini-Hochberg method [47] to correct for multiple comparisons for all post-hoc tests unless stated otherwise. Fluorescence changes from baseline were assessed against zero using one-sided Wilcoxon signed-rank tests. Differences in neuronal and astrocytic responses across stimulation intensities, including response peak, fractions of responsive ROIs, and recruitment dynamics, were evaluated using Kruskal-Wallis tests, followed by pairwise Mann-Whitney post-hoc tests when appropriate. Relationships between response peak and stimulation intensity were examined using Pearson correlation for z-scored response peaks and Spearman correlation for spatial measures such as ROI distance. Distributions of responsive versus non-responsive ROIs, including comparisons between neuronal and astrocytic layers and between ROIs containing neuronal soma versus those without, were tested using chi-squared analysis. Pharmacological controls with TTX and comparisons across stimulation intensities were analyzed using the same statistical frameworks as baseline tests.

*Code is available in an accessible repository:* https://github.com/GrunyD/Neuron_astrocyte_stimulation_analysis

## Results

### Astrocytes respond to stimulation amplitude levels as low as those activating neurons

To investigate astrocytic recruitment by ICMS, we performed stimulations *in vivo* anaesthetized mice, carefully monitored to ensure physiological parameters within healthy ranges during the entire experiment (Fig. 1A). We performed dual-color two-photon calcium imaging with co-labeled astrocytic and neuronal indicators in mouse visual cortex, allowing direct comparison of astrocytic and neuronal responses to the stimulations (Fig. 1B and C). We used variable stimulation intensities, during an invariable ICMS protocol (Fig. 1D). Figure 1E shows the averaged neuronal Ca^2+^ response amplitudes across the entire field of view during stimulation intensities ranging from 10 to 70µA. In the same imaging field, astrocytic Ca^2+^ responses (Fig. 1F) were also robust and significant even though our stimulation duration was 0.5ms, which is transient compared to the previous study [27]. Notably, even at the lowest stimulation intensity tested (10µA), both neurons and astrocytes exhibited reliable responses, although astrocyte amplitudes were lower than neuronal ones. Because spontaneous Ca^2+^ activity in the absence of stimulation shows no covarying fluctuations between neuronal and astrocytic channels (supplementary Figure1), the astrocyte responses observed at low stimulation intensities are unlikely to be attributable to channel bleaching. Across all stimulation levels, responses remained significantly elevated relative to baseline and were consistent across animals for both cell types (Fig. 1E-F, right, *p*<0.05). These results demonstrate that astrocytes can be effectively recruited *in vivo* at stimulation intensities comparable to those activating nearby neurons, in contrast to previous reports suggesting minimal astrocytic responsiveness at low-intensity ICMS [23,27]. Moreover, these findings indicate that they are an active component of the ICMS-evoked network alongside neurons, exhibiting dynamics similar to those observed during *in vivo* sensory stimulation [48–50].

### Neuronal activation depends on both stimulus intensity and proximity to the electrode, while astrocyte responses are intensity-linked but lack spatial gradients

As astrocytic morphology is highly complex, with fine processes that make region-specific quantification challenging [32], we employed a grid-based analysis. This approach disregards detailed morphological features and instead uniformly samples across all astrocytic anatomical regions, enabling quantification of spatiotemporal variability in astrocytic and neuronal responses across repeated stimulations. To avoid imprecise response measurements caused by repetitive stimulation trials and randomly drifting baseline, we further applied a baseline subtraction to each ROI (Fig. 2B). We then used a threshold criterion (see *Methods* for details) to classify each ROI as responsive or non-responsive to stimulation. Although we observed significant astrocytic responses in Figure 1F where the signals were averaged across the full field of view, we found that when we divided the imaged field into grids, astrocytes exhibited lower peak responses (Fig. 2C, lower) compared to neurons (Fig. 2C, upper). Pooled analysis across all stimulation-responsive ROIs revealed a gradual and systematic increase in neuronal peak responses with increasing stimulation intensity, reflected by a significant positive correlation (Fig. 2D, left, pearson *r*=0.519, *p*<10^-15^). In contrast, astrocytic peak responses exhibited weaker modulation by stimulation intensity, showing a low correlation with increased stimulation intensity compared to neurons (Fig. 2D, right, pearson *r*=0.112, *p*<10^-15^).

**Figure 2.**
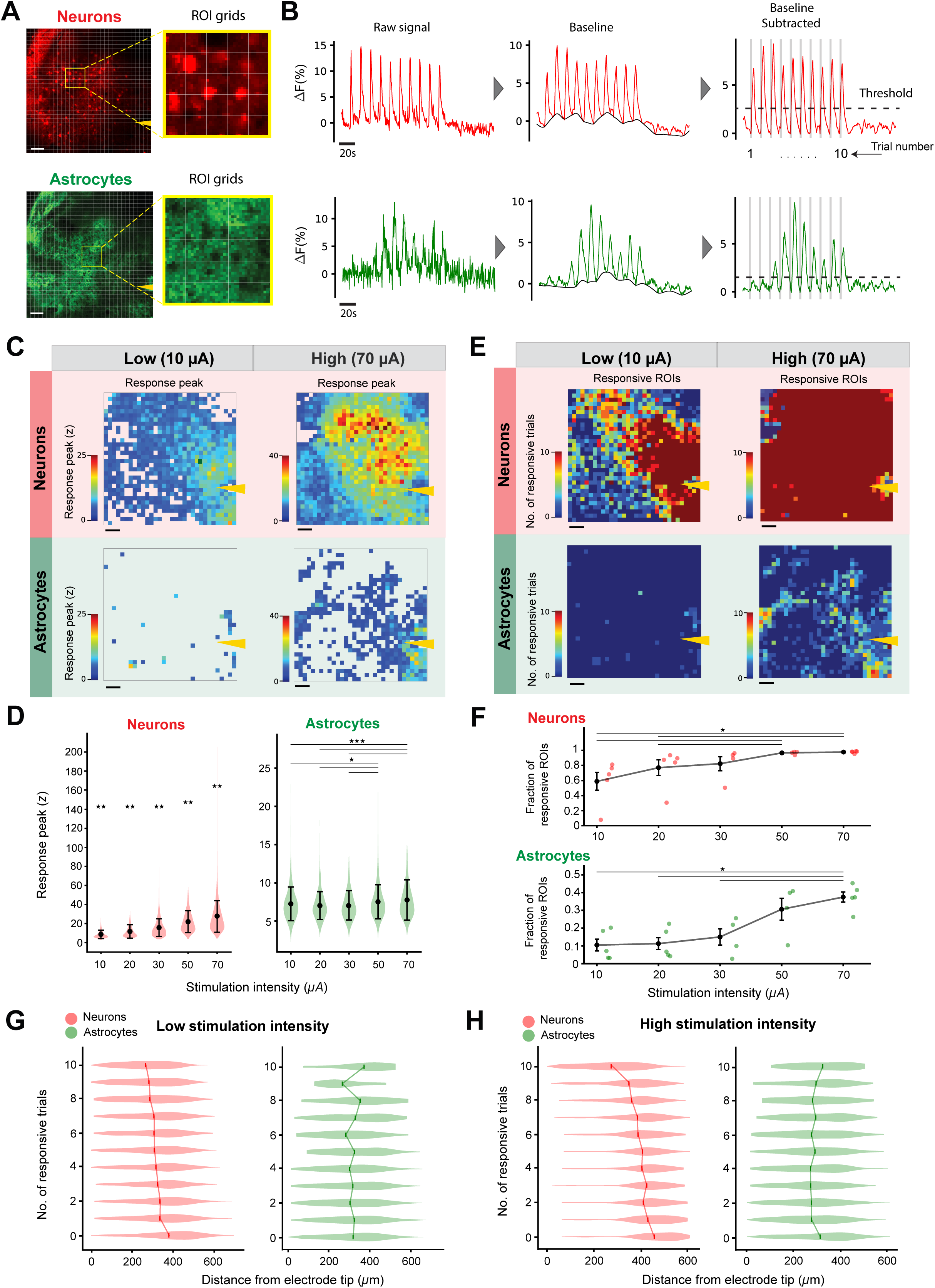
Grid-based analysis reveals strong distance-dependent neuronal activation, contrasted with weak distance dependency in astrocytes. **A.** Representative imaging fields showing the neuronal channel (top) and astrocytic channel (bottom), each divided into ROI grids for spatial analysis. Scale bar, 50µm. **B**. Example traces from one representative ROI. Top: Raw neuronal Ca2+ signal (left), baseline estimation (black trace, middle), and baseline-subtracted trace (right) used for threshold-based response detection (dashed horizontal line). Gray bars indicate stimulation timing; trial numbers (1-10) are shown below. Bottom: Same convention as above, but for the astrocytic signal. **C**. Changes in response peak across ROI grids for neurons (top) and astrocytes (bottom) under low-(10µA) and high-amplitude (70µA) stimulation intensity. Scale bar, 50µm. **D**. Pooled response peaks across all ROIs as a function of stimulation intensity for neuronal ROIs (left) and astrocytic ROIs (right). Error bars represent mean +/-SD. Lines above data points denote paired comparisons tested with the Mann-Whitney test. \**p*<0.05, \*\**p*<0.01,\*\*\**p*<0.001. **E**. Same format as in **C**, but showing the number of responsive trials for each responsive ROI (see *Methods* for the definition of “responsive ROI”). **F**. Same convention as in D, but responses were pooled within individual mice, and the fraction of responsive ROIs was plotted as a function of stimulation intensity (each dot represents one mouse). Neuronal responses are shown above; astrocytic responses are shown below. Kruskal-Wallis test, \**p*<0.05. **G**. The number of responsive trials is plotted as a function of the ROIs distance from electrode tip. Low stimulation intensity conditions (≤30µA). A line connects the mean number of responsive trials across distance. **H.** Same convention as G, but high stimulation intensity conditions (≥50µA).

Despite this variation in amplitude scaling, the fraction of responsive ROIs in both neurons and astrocytes differed significantly across stimulation intensities (Fig. 2F, *p*<0.001). Notably, pairwise comparisons revealed a marked increase in the fraction of responsive astrocytic ROIs between 30µA and 50µA (Fig. 2F, lower panel), indicating that higher stimulation intensities recruit a larger number of astrocytic territories even though their individual peak responses do not scale as strongly as those of neurons.

To further characterize the spatial organization of the responses, we quantified ROI responsiveness as a function of distance from the electrode tip. Neuronal ROIs located closer to the electrode tip showed a higher number of responsive trials, which were defined as those with a response amplitude above a fixed threshold (Fig. 2B, example trace with the dashed horizontal line). This finding is consistent with the expected effect of the local electric field, which likely influences nearby neurons more strongly [51,52]. The relationship was reflected in significant distance-responsiveness correlations under both low- and high-intensity stimulation (Fig. 2G, H, Left: Spearman *r*=-0.32, *p*<0.001 at low stimulation intensity; *r*=-0.36, *p*<0.001 at high stimulation intensity). In contrast, astrocytic ROIs showed more spatially dispersed and less distance-dependent activation patterns (Fig. 2G, H, Right; Spearman *r*=-0.007, *p*<0.001 at low stimulation intensity; *r*=-0.085, *p*<0.001 at high stimulation intensity). These results indicate that neuronal responses to ICMS scale systematically with both stimulation intensity and proximity to the electrode, whereas astrocytic responses increase in overall recruitment at higher amplitudes but lack a strong spatial bias toward the electrode tip.

### Astrocytic responses show greater temporal variability than neurons across repetitive stimulation

Next, we asked how robustly neurons and astrocytes respond during repetitive stimulation trials. As shown in Figure 2B, the representative traces showed that astrocytic response was less uniform across the 10 stimulation trials compared to neuronal responses (Fig. 2B, lower green trace). To assess the generality of this phenomenon across ROIs, we identified both the first responsive peak and the highest responsive peak over all trials. Consistent with the representative data, most neuronal ROIs exhibited their first responsive peak during the first trial, whereas astrocytic ROIs showed a more dispersed distribution for both the first and highest responsive peaks (Fig. 3A). With regards to the highest peak, this temporal dispersion was most pronounced at low stimulation intensities. At higher stimulation intensities, astrocytic responses tended to occur earlier in the trial sequence, with both the first and highest peaks appearing with the first stimulations (Fig. 3A).

**Figure 3.**
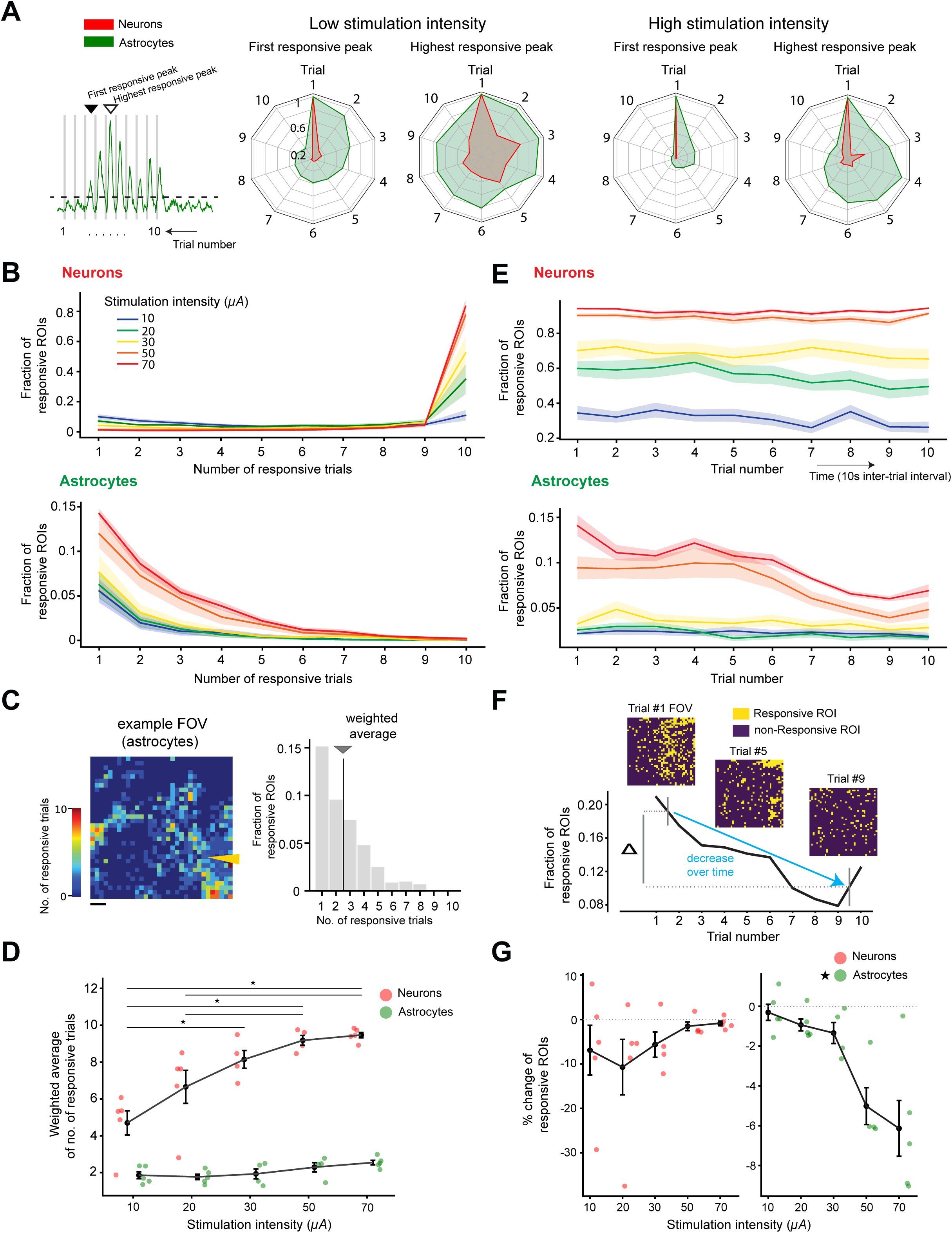
Astrocytic responses are temporally variable and attenuate at high stimulation intensity compared with stable neuronal responses. **A.** *Left*: Example trace from Figure 2B illustrating the “first responsive peak” (open triangle) and the “highest responsive peak” (filled triangle). *Right:* Spider plots summarizing the temporal distribution of responses. The first two plots show, respectively, the fraction of first responsive peaks and the fraction of highest responsive peaks at low stimulation intensities (≤30uA), each normalized to the maximum response observed across 10 trials. The two rightmost spider plots show the corresponding distributions at high stimulation intensities (≥50µA). **B.** Fraction of responsive ROIs plotted as a function of the number of responsive trials for neurons (top) and astrocytes (bottom). **C**. Example field of view showing astrocytic ROIs color-coded by the number of responsive trials (left). The corresponding histogram (right) shows the fraction of responsive ROIs across trial counts. The dark gray line and arrow indicate the weighted average number of responsive trials. Scale bar, 50µm. **D**. Weighted averages from (C) plotted for each mouse (one dot= one mouse data) across stimulation intensities. Lines indicate pairwise comparison (Mann-Whitney test, \**p*<0.05). **E**. Fraction of responsive ROIs plotted across actual trial numbers for neurons (top) and astrocytic (bottom). The color scheme corresponds to panel (B). Each trial was separated by a 10 second inter-trial interval. **F**. Example illustrating the decline in the fraction of responsive ROIs across trials (y-axis). Insets show responsive (yellow) and nonresponsive (dark purple) astrocytic ROIs at the 1st, 5th, and 9th trials. These trial-wise changes were quantified for group-level comparison. **G**. Percent change in the fraction of responsive ROIs across trials plotted as a function of stimulation intensity.

Further, the number of neuronal responsive ROIs increased systematically with stimulation intensity (10µA-70µA), reflected by a greater proportion of ROIs responding across more trials at higher currents (Fig. 3B, upper). In contrast, astrocytic responses were skewed toward ROIs that responded in a limited number of trials, a sparsity that was evident at all tested stimulation intensities (Fig. 3B, lower). To quantify these trends, we calculated the weighted average number of responsive trials for each responsive ROI (see *Methods* for details; Fig. 3C). Across all animals, the number of neuronal responsive trials increased with stimulation intensity, reaching an average of 10/10 response/trials with the highest stimulation intensity (Fig. 3D, Kruskal-Wallis test, *p*<0.01). In contrast, astrocytic ROIs showed relatively stable trial counts across intensities, on average 2/10 (Fig. 3D, Kruskal-Wallis test, *p*>0.05), indicating that higher stimulation intensities recruit more astrocytic territories but do not increase the reliability with which individual astrocytic ROIs respond. Together, these results suggest that astrocytes exhibit greater temporal variability than neurons during repetitive stimulation, reflecting distinct modes of temporal integration in glial versus neuronal populations under the same ICMS conditions.

### Astrocytic desensitization to repeated high stimulation intensity

One of the notable findings was that astrocytic responsiveness declined across repeated trials at higher stimulation intensities. Starting at 50µA, the fraction of responsive astrocytic ROIs significantly decreased over the last five trials (Fig. 3E, lower panel). To quantify this effect, we measured the change in the fraction of responsive ROIs across trials (Fig. 3F). Astrocytic responses showed a slight decline at lower stimulation intensities (<30µA), but a pronounced reduction emerged at 50µA and above (Fig. 3G, right panel, Kruskal-Wallis test, *p*<0.05). Interestingly, this trial-to-trial decrease was absent in neurons, whose response fractions remained stable across both trials and stimulation intensities (Fig. 3E, top; G, left panel, Kruskal-Wallis test, *p*>0.05).

Neuronal stimulation triggers glutamate release, which subsequently evokes Ca^2+^ elevations in neighboring astrocytes [53], and astrocytic Ca^2+^ signals can, in turn, modulate neuronal activities via gliotransmitter release. Under high-frequency repetitive ICMS, glutamate-associated astrocytic calcium accumulation may introduce slow baseline fluctuations. Because our preprocessing pipeline includes baseline subtraction, it’s important to determine whether increased astrocytic baseline activity could artificially contribute to the apparent reduction in astrocytic responsiveness at high stimulation intensities observed in Figure 3G. We found that both neuronal and astrocytic signals showed elevated baseline levels at high stimulation intensities compared to that of low stimulation intensities (Supplementary Figure 2A, *p*< 0.001). Neurons showed even higher baseline elevation compared to astrocytes at high stimulation intensity. In addition, for both cell types, baseline activity was significantly lower during the last five trials compared to the first five trials (Supplementary Figure 2B, *p*<0.001). These results indicate that the observed reduction in astrocytic responsiveness at high stimulation intensities cannot be explained simply by shifts in baseline fluorescence, thereby excluding the possibility that the effect is due to a preprocessing artifact. Overall, astrocytic responses desensitized with repeated, high-intensity stimulation, while neuronal responses remained consistent across trials despite an accompanying rise in baseline activity.

### Astrocytes show increased responsiveness near active neuronal somata

So far, we have independently examined the responsiveness of neuronal and astrocytic ROIs in the whole field of view or in grids. Next, we further investigated whether astrocytic responsiveness is influenced by neuronal activity by analyzing the relationship between responsive neuronal and astrocytic ROIs (Fig. 4A illustrates the method). Interestingly, ROIs where the astrocytes were responsive exhibited significantly higher neuronal peak response intensity, compared to ROIs in which the astrocytes were nonresponsive (Fig. 4B). Notably, when neuronal ROIs responded consistently across all 10 trials, the corresponding astrocytic ROIs showed both the highest fraction of responsiveness and the greatest peak intensity (Fig. 4C, D). A similar trend was found in both low and high stimulation intensities (Supplementary Figure 3A, B).

**Figure 4.**
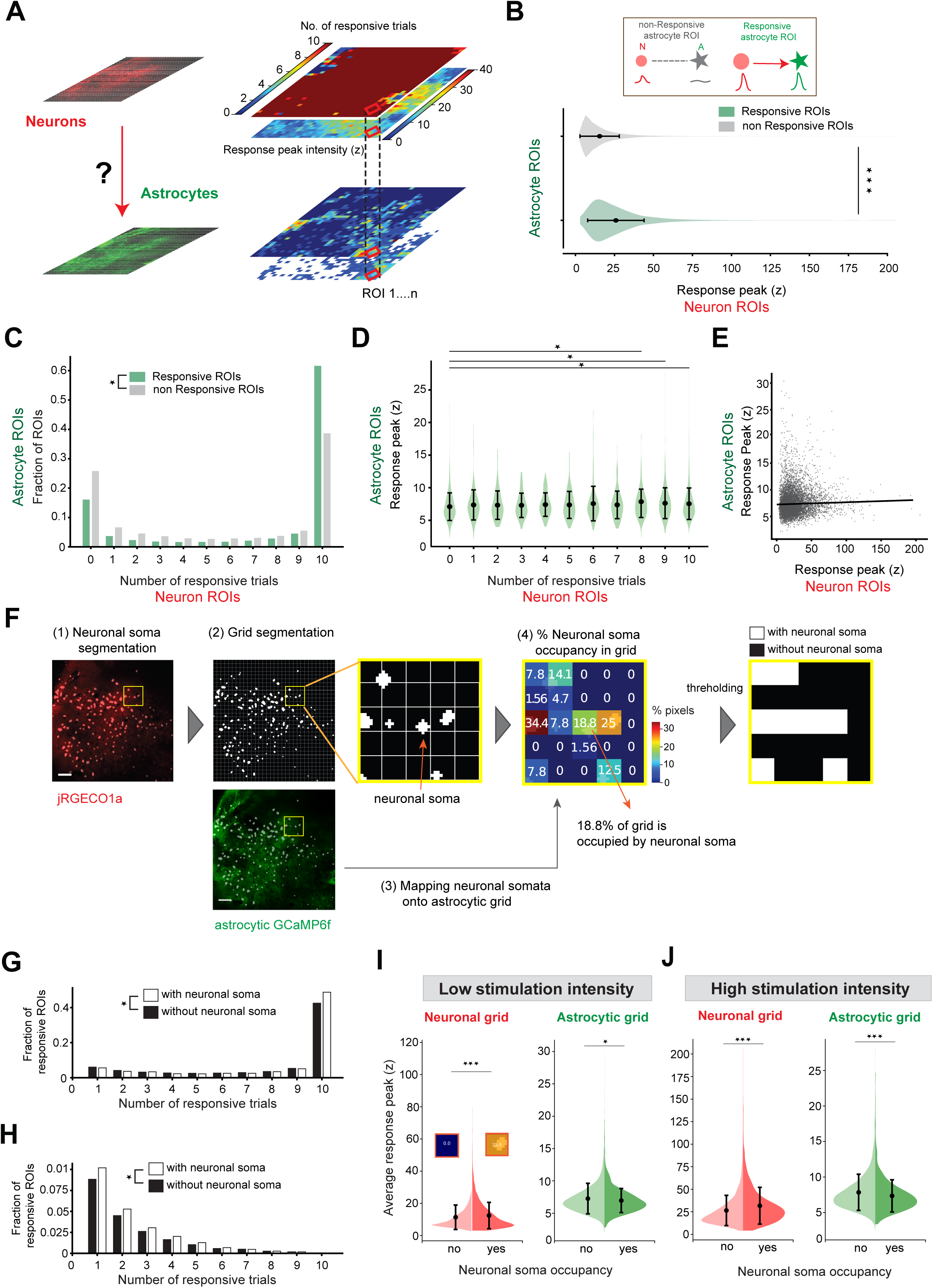
Astrocytic responsiveness increases with strong neuronal activation and is higher in ROIs containing neuronal soma. **A.** Schematic illustration of the method: responsive neuronal ROIs were identified and spatially overlaid to extract corresponding ROIs from the astrocytic channel. **B.** Violin plots comparing astrocytic ROIs categorized as responsive (green) or nonresponsive (gray), plotted against the peak response of their paired neuronal ROIs. (Mann-Whitney test, \*\*\**p*<0.001). **C**. Fraction of responsive (green) and nonresponsive (gray) astrocytic ROIs as a function of the number of responsive trials. (Chi-square test, \**p*<0.001). **D**. Astrocytic peak responses plotted against neuronal responsive trial count, following the same color convention as in C. Pairwise comparisons are indicated by lines above violin plots (Mann-Whitney test, \**p*<0.05). **E**. Control analysis showing no correlation between astrocytic (y-axis) and neuronal (x-axis) response peaks. Black line indicates linear regression (pearson *r* = 0.0078, *p*< 0.001). **F.** Illustration of neuronal somata segmentation (1), projection to astrocytic grid (2), and quantification of neuronal soma occupancy in each ROI with thresholding (3-4; see *Methods* for details). Scale bar, 50µm. **G.** Fraction of responsive ROIs as a function of the number of responsive trials in neuronal grid. Open squares indicate grids containing neuronal somata: closed squares indicate grids without neuronal somata. Chi-square test, \**p*<0.01**. H.** Same as G but astrocytic grid. **I-J**. Violin plots comparing astrocytic peak responses between ROIs containing neuronal somata and those without. Low stimulation intensity (I, ≤30μA) and high stimulation intensity (J, ≥ 50μA). Mann-Whitney test, \**p*<0.05, \*\*\**p*<0.001.

As the aforementioned results indicated that the number of responsive trials in neurons are well correlated with astrocytic responsivity, we further examined whether there was a correlation between peak intensities of neuronal and astrocytic responses in all ROIs. The analysis revealed no significant correlation (Fig. 4E), which eliminates any concern that this coupling was due to imperfect separation of the emission from the two fluorophores leading to detection of mixed signals.

We next asked whether astrocytic responsiveness depended on presence of neuronal soma in close proximity. While astrocytic activation around processes reflects synaptic transmission, activation near somas might be dependent on the recruitment of that entire specific neuron in the network and its summed response to the stimulation. To address this, we segmented neuronal somata within each ROI and projected their locations onto the astrocyte channel. We then calculated the percentage of neuronal soma occupancy per ROI and classified ROIs as either containing somata or not, using a 5% occupancy threshold (Fig. 4F). Intriguingly, ROIs containing neuronal somata exhibited a higher fraction of responsive ROIs in both neuronal and astrocytic populations (Fig. 4G-H), suggesting that astrocytic responsiveness depends on nearness to neuronal soma. This said, Figure 4 I and J show that astrocytic responses are largest further away from the soma. Since the density of synapses is expected to be largest in ROIs that do not contain a neuronal somata, this indicates that the increase in astrocytic Ca^2+^ levels depends on synaptic transmission.

### Neuronal spiking drives astrocytic responses

To this point, our analysis revealed distinct spatiotemporal dynamics of astrocytic responses to ICMS compared to neuronal activity. The variable and spatially heterogeneous astrocytic responses could indicate a direct recruitment by the stimulation circumventing the neurons. To investigate their origin, we topically applied tetrodotoxin (TTX) to block actional potentials (Fig. 5A) and analyzed responsive ROIs for both cell types using the same protocol (Fig. 5C). Representative traces (Fig. 5B) and population analyses (Fig. 5C, D) showed that over 98% of responsive neuronal ROIs were abolished following TTX application, with astrocytic ROIs exhibiting a similar suppression. Only a small fraction of ROIs remained responsive, typically appearing in just a few of the 10 trials (Fig. 5D). At the population level, neither astrocytes nor neurons showed responses that differed from baseline (Fig. 5E, *t*-test against zero, *p*>0.05). These results indicate that although astrocytes differ from neurons in both response consistency and their dependence on stimulation intensity, they depend on neuronal activation. The elimination of neuronal responses by TTX confirms that astrocytic activation is not directly evoked by electrical stimulation but instead arises secondarily through neuronal activity.

**Figure 5.**
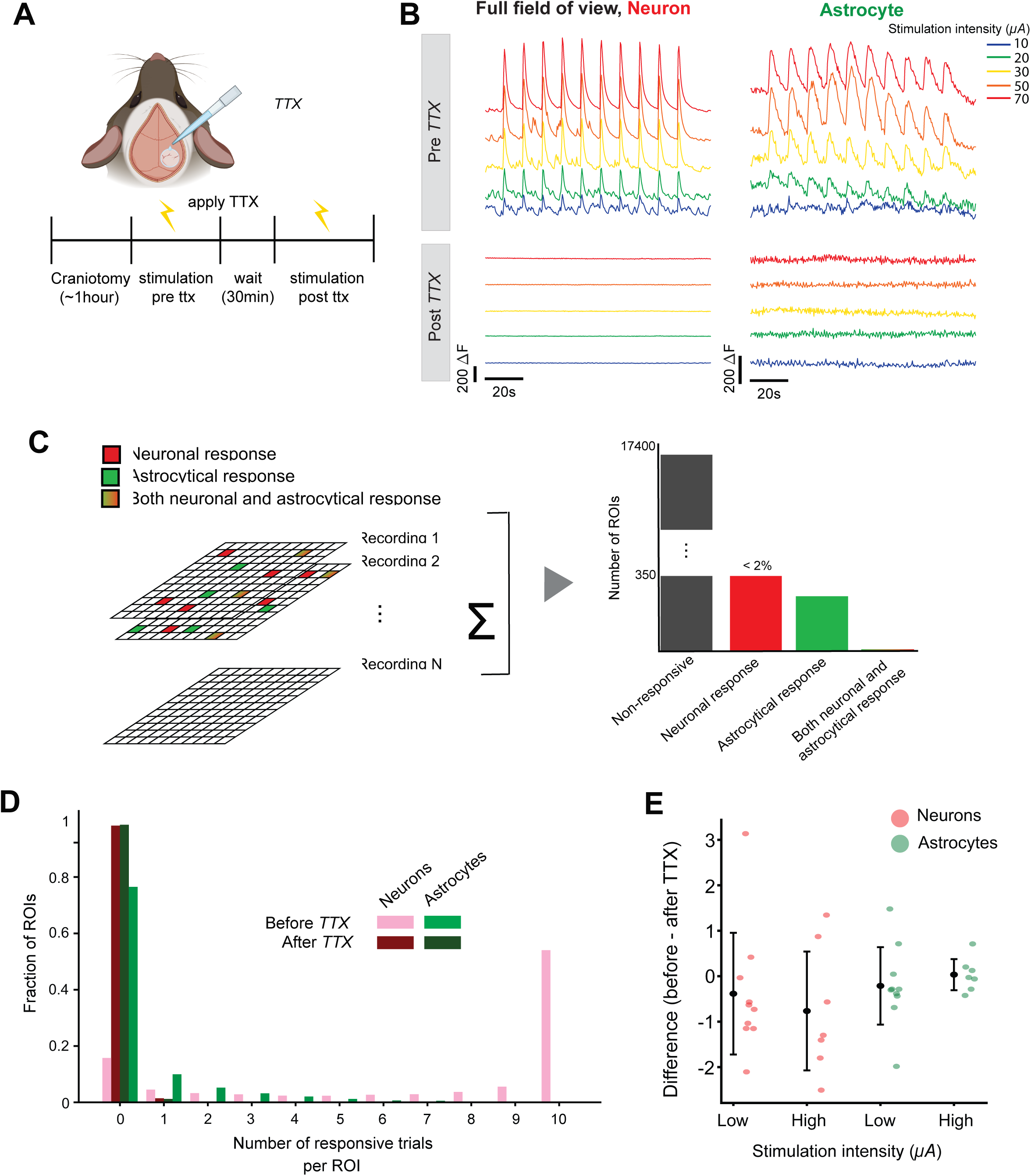
Both neuronal and astrocytic responses are abolished by tetrodotoxin. **A.** Schematic of the TTX experiment. **B** Representative traces of neuronal and astrocytic responses before and after TTX application across different stimulation intensities. **C**. Grid-based analysis of neuronal and astrocytic ROIs responsive to stimulation, with histogram showing the percentage of responsive ROIs after TTX. **D**. Fraction of ROIs plotted against the number of responsive trials: before TTX, (neurons, red; astrocytes, green) and after TTX (neurons, dark red; astrocytes, dark green). **E**. Population-level differences (one dot=one mouse) in response peak before and after TTX during low (≤30μA) and high (≥50μA) stimulation intensities.

## Discussion

### Summary

We found that astrocytes are robustly recruited during ICMS-even at current as low as 10μA, yet their responses differ from neuronal responses. They also show weaker spatial dependence on the electrode tip compared to neurons, whose activation strongly correlates with proximity to the stimulation site. Neuronal responses were consistent across repeated trials, whereas astrocytic activity exhibited greater temporal variability. Notably, while the number of responsive trials in astrocytes remained relatively constant across stimulation intensities, the spatial extent of their responsive territories expanded with increasing intensity. Astrocytic signals showed a form of desensitization, with response peak diminishing in later trials - a phenomenon absent in neurons. Despite their spatially diffuse and temporally variable activation patterns, astrocytic responsiveness was strongly dependent on neuronal activity. Our analyses reveal both synchronous and asynchronous modes of neuron-astrocyte interaction under identical stimulation conditions, highlighting astrocytes as an integral yet distinct component of electrically evoked cortical dynamics.

### Integrative and network-based mechanisms of astrocytic calcium signaling underlying ICMS engagement

Astrocytic Ca^2+^ activity is highly heterogeneous, spanning rapid microdomain transients to slower, spatially extended events lasting seconds to minutes [10,44,54–56]. This diversity arises from subcellular inputs mediated largely by neurotransmitter activation of astrocytic G-protein-coupled receptors (GPCRs) [57], including glutamatergic, dopamine, acetycholine and γ-aminobutyric acid (GABA) pathways [40]. The signaling routes produce distinct temporal profiles, for example, glutamatergic inputs evoke rapid and robust Ca^2+^ elevation, whereas GABA_b_-mediated receptor activation generates delayed and prolonged responses [58]. These GPCR-mediated signals relay on intracellular inositol (1,4,5)-triphosphate receptor (IP3R)-dependent Ca^2+^ release from the endoplasmic recticulum (ER) [59] rather than fast, channel-mediated Ca^2+^ influx typical of neurons [60,61], resulting in slower onset, decay, and greater temporal variability. Recently, it has been shown that whereas the largest of the astrocytic Ca^2+^ responses are dependent on the IP3R2 receptor subtype [62], astrocytic responses at synapses rely on the IP3R1 subtype. Thus astrocytic Ca^2+^ recruitment differs spatially with regards to synaptic nearness [63]. Astrocytes are also interconnected through gap junctions [64,65] and embedded within fine perisynaptic microdomains [66–68], enabling locally triggered Ca^2+^ elevations to propagate across extensive territories and integrate activity from diverse neuronal sources. Thus, astrocytic calcium signaling reflects an integrative and network-based process [44,69] shaped by both local receptor activation and multicellular propagation. These properties position astrocytes to respond to ICMS not through direct electrical activation but via the downstream consequences of stimulation-evoked neuronal activity, as supported by our TTX results and previous work [27]. Therefore, the integrative and propagative nature of astrocytic signaling provides a mechanistic framework for understanding the spatially extended and temporally heterogeneous astrocytic responses observed in our experiments.

### Spatial and temporal characteristics of astrocyte recruitment during ICMS

Our grid-based analysis approach, which captures Ca^2+^ activity across all structures within a defined spatial grid, enabled us to characterize patterns consistent with the mechanistic principles. The spatially diverse astrocytic responses we observed - showing only weak dependence on the distance from the stimulating electrode-support the view that astrocytes are not directly activated by ICMS but are instead driven by neurotransmitter-mediated signaling, including glutamatergic and nonadrenergic pathways [27]. That astrocytic response can occur at a distance from the triggering neurotransmitter and without obvious waves of Ca^2+^ activity spreading through the syncytium was shown in a recent uncaging-experiment of both glutamate and GABA [58]. This aligns with prior work demonstrating that electrical stimulation involves synaptic transmission of neurons, mainly by the glutamate pathway [70–72], which can diffuse through the extracellular space to engage astrocytic GPCRs. Although neurons distal to the electrode tip can be recruited, as shown in previous work and in our results [23,46,73–75], leading to additional neurotransmitter release, our finding that astrocytic activation was only weakly correlated with distance from the stimulating electrode indicates that astrocytes are less spatially constrained than neurons during ICMS. It is also worth noting that the spatial extent of astrocyte recruitment scaled with stimulation intensity in parallel with the intensity-dependent expansion of neuronal activation. This coupling suggests that astrocytic engagement largely reflects the downstream consequences of increased neuronal recruitment at higher intensities, superimposed on the intrinsic integrative properties of the astrocytic network.

Temporally, increasing stimulation intensity did not increase the number of trials in astrocytic responses within a grid (Fig. 3D), in contrast to neurons, which remained reliably activated across all repetitions. This temporal dispersion is particularly evident in the distribution of the largest response peaks across 10 repeated stimulations at both low and high intensities (Fig. 3A). Such variability is well explained by the slower GPCR-IP3R-dependent Ca^2+^ dynamics of astrocytes [76,77], which involve ER-mediated release that can exhibit partial Ca^2+^ accumulation or refractory periods. These intrinsic kinetics, combined with rapid glutamatergic activation [40,58] across diverse astrocytic microstructures, likely limit astrocytes’ ability to respond consistently during minute-scale repetitive ICMS.

Despite slight differences in stimulation parameters, our findings align with Ma et al., (2021)[27], who similarly reported that astrocytic Ca^2+^ elevation could not be sustained during prolonged electrical stimulation, in contrast to the stable responses observed in neurons. Although their study used single continuous stimulation lasting up to 30 seconds, whereas we applied brief (0.5s) pulses repeated across 10 trials, both studies converge on the observation that astrocytic responsiveness diminishes with time progression. In our experiment, this phenomenon was particularly pronounced at higher stimulation intensities: astrocytic responsiveness progressively declined at later half of the stimulation sequence, corresponding to the period after approximately one minute of repeated ICMS (Fig. 3E), while neuronal responses remained stable and robust across all trials and intensities. We further found that this decline in astrocytic responsiveness at high intensities coincided with an elevation in baseline Ca^2+^ level in both neurons and astrocytes. Although neuronal evoked responses were preserved within our observation window, the longer-term implications of this baseline shift remain unknown. Astrocyte-neuron communication is a bidirectional and integrative regulatory system [39], such that neurons initiate astrocytic activation, and astrocytes, in turn, can modulate neuronal GPCR-dependent Ca^2+^ signaling via gliotransmitters including glutamate, D-Serine, ATP [78–81], influence synaptic plasticity [82], and are involved in cognitive processing [13]. Whether the Ca^2+^ accumulation observed here ultimately alters neuronal function or affects network stability-either beneficially or detrimentally under ICMS, is an open question for further investigation.

### Implications for neuroprosthetic and brain-machine interface design

While glial responses to device implantation have been extensively studied [83–85], astrocytic engagement during ICMS has important implications for neuroprosthetic performance, as well as emerging therapeutic applications [86–88]. Because astrocytes integrate neuronal activity over broader spatial and temporal windows, their recruitment, even indirectly via neurotransmitter signaling, may influence how stimulation-evoked signals propagate within cortical circuits, beyond the stimulated region [87]. Astrocytic Ca^2+^ dynamics can modulate synaptic efficacy, local excitability, neurotransmitter clearance, and neuromodulator release, all of which are relevant for encoding artificial percepts or maintaining stable long-term stimulation performance. The temporally heterogeneous nature of astrocytic activity observed in our study, even with low current, suggests that glial networks may contribute to shaping the perceptual spread, intensity tuning, and neural adaptation phenomena that are important in ICMS-based sensory restoration paradigms.

Moreover, our observation that repeated ICMS produces intensity-dependent baseline Ca^2+^ elevation in both neurons and astrocytes raises considerations for chronic neuroprosthetic use. While neuronal responses remained robust over the duration of our experiments, astrocytic desensitization or adaptation and cumulative calcium load could influence network excitability, plasticity, or inflammatory signaling over longer time scales even though this awaits further investigation. Together, our findings highlight the importance of considering both neuronal and astrocytic dynamics when designing ICMS protocols for optimizing stimulation parameters to balance efficient neuronal recruitment with controlled astrocytic engagement, which may help improve perceptual fidelity, and support long-term stability of implanted systems.

## Acknowledgments

David Grundfest and Kayeon Kim contributed equally to this work.

C.C acknowledges support from the Lundbeck Foundation (#R345-2020-1440, #R436-2023-1125, #R392-2018-2266, and #R345-2020-1440), the Independent Research Fund Denmark (#1133-00016B and #1030-00374A), the Novo Nordisk Foundation (#0064289, #0092323, #117272, #14CC0001), Læge Sofus Carl Emil Friis og Hustru Olga Doris Friis’ Legat, Dagmar Marshalls Fond, Familien Hede Nielsens Fond, Hørslev fonden, Oda og Hans Svenningsens Fond, and Helsefonden. B.L acknowledges support from the Independent Research Fund Denmark (#1030-00374B), Læge Sofus Carl Emil Friis og Hustru Olga Doris Friis’ Legat, Dagmar Marshalls Fond, Familien Hede Nielsens Fond, Hørslev fonden, and Oda og Hans Svenningsens Fond. K.Kim acknowledges support from Hørslev fonden, Dagmar Marshalls Fond, and Oda og Hans Svenningsens Fond. We used AI language tools for grammar checking and language refinement during manuscript preparation. Some images were created using Biorender.com under license.

---

**Supplementary Figure 1.**
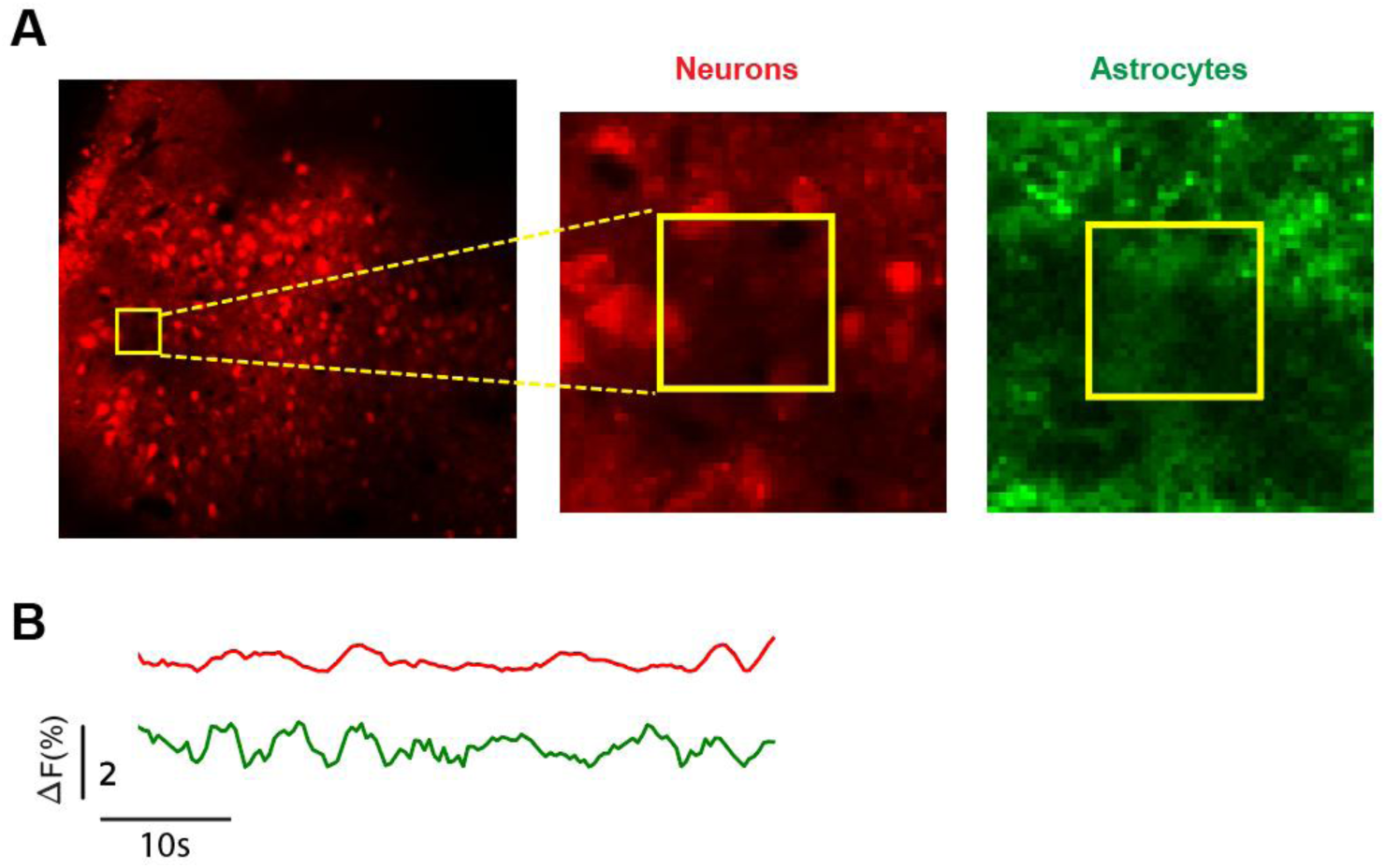
Spontaneous Ca^2+^ activity of neuronal and astrocytic grids to assess potential channel bleaching effects. **A.** A representative field of view (FOV) illustrating the grid-based segmentation used for analysis. The yellow grid corresponds to a group of nine adjacent subregions (small squares) to check background Ca^2+^ fluorescence. **B**. Average Ca^2+^ fluorescence traces recorded during non-stimulation periods from the yellow grid, neurons (red) and astrocytes (green). Traces show lack of co-varying fluctuation, indicating no channel bleaching under the experimental conditions.

**Supplementary Figure 2.**
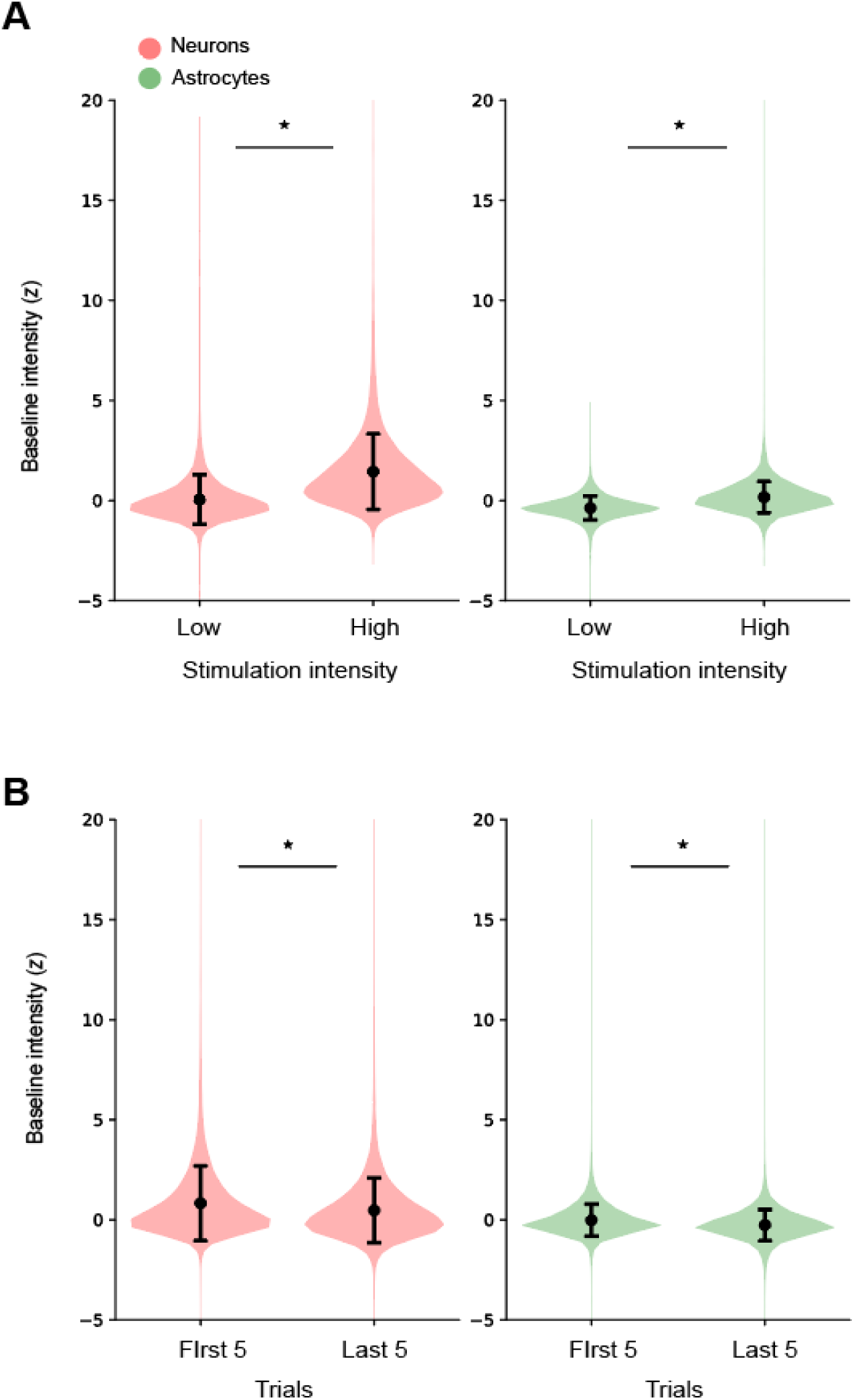
Elevated baselines at high stimulation current and trial-dependent declines coincide with reduced astrocytic responsiveness. **A.** Baseline intensity of neuronal (left) and astrocytic (right) responses at low (≤30μA) and high (≥50μA) stimulation intensities. Mann-Whitney test, \**p* < 0.001. **B**. Same convention as A, but baseline intensity of neuronal (left) and astrocytic (right) responses at the first five and the last five trials.

**Supplementary Figure 3.**
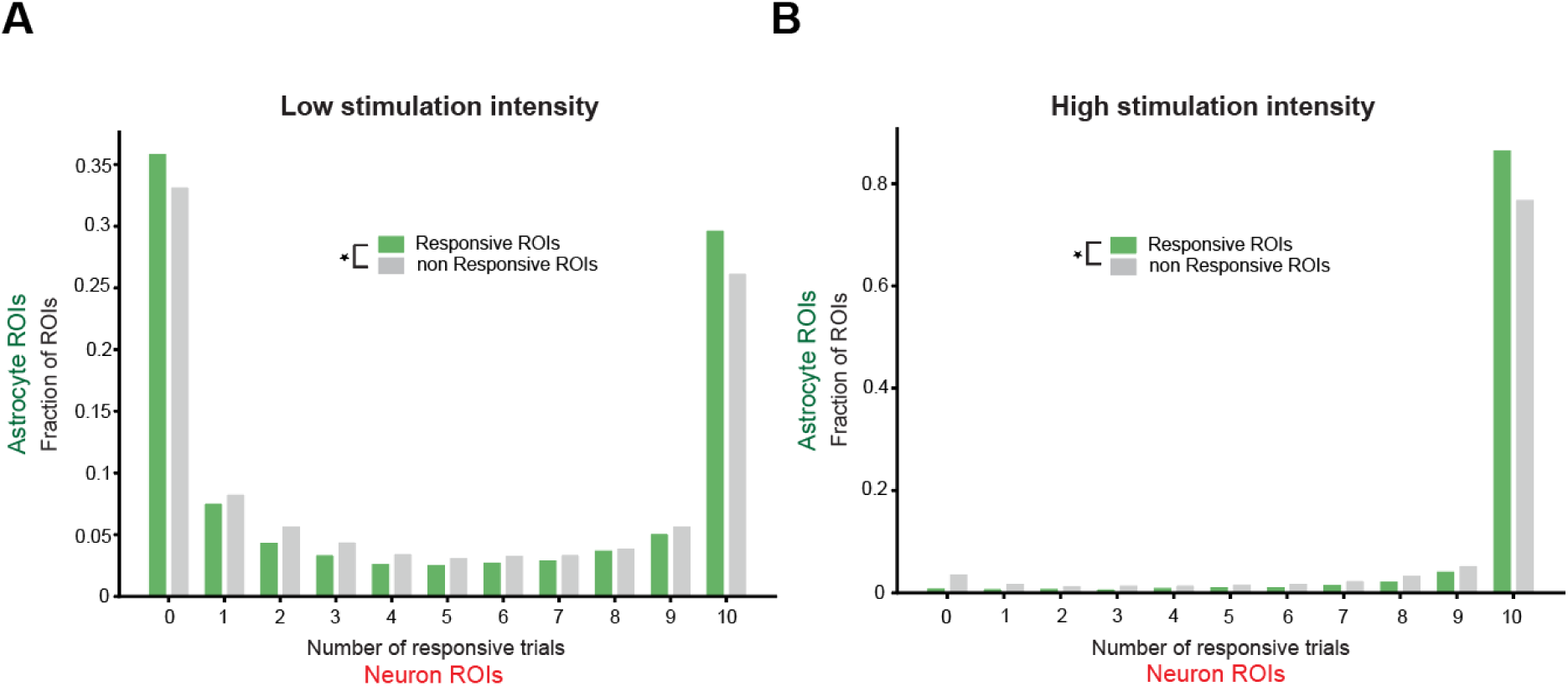
Astrocytic responsiveness is related to the neuronal soma proximity in both low and high stimulation intensities. **A-B**. Fraction of responsive (green) and nonresponsive (gray) astrocytic ROIs plotted against the number of responsive trials, separated by stimulation amplitude: low (A, ≤30μA) and high (B, ≥50 μA). Chi-square test, \**p* < 0.001.

